# Cryo-EM-guided engineering of T-box-tRNA modules with enhanced selectivity and sensitivity in translational regulation

**DOI:** 10.1101/2023.02.28.530422

**Authors:** Xinyu Jia, Chong Zhang, Bingnan Luo, Jane K. Frandsen, Andrew M. Watkins, Kaiyu Li, Minghua Zhang, Xiawei Wei, Yang Yang, Tina M. Henkin, Zhaoming Su

## Abstract

Riboswitches are non-coding RNA elements that play vital roles in regulating gene expression. Their specific ligand-dependent structural reorganization facilitates their use as templates for design of engineered RNA switches for therapeutics, nanotechnology and synthetic biology. T-box riboswitches bind tRNAs to sense aminoacylation and control gene expression via transcription attenuation or translation inhibition. Here we determine the cryo-EM structure of the wild-type *Mycobacterium smegmatis ileS* T-box in complex with its cognate tRNA^Ile^. This structure shows a very flexible antisequestrator region that tolerates both 3’-OH and 2’,3’-cyclic phosphate modification at the 3’ end of tRNA^Ile^. Elongation of one helical turn (11-base pair) in both the tRNA acceptor arm and T-box Stem III maintains T-box-tRNA complex formation and increases the selectivity for tRNA 3’ end modification. Moreover, elongation of Stem III results in ∼6-fold tighter binding to tRNA, which leads to increased sensitivity of downstream translational regulation indicated by precedent translation. Our results demonstrate that cryo-EM can guide RNA engineering to design improved riboswitch modules for translational regulation, and potentially a variety of additional functions.

## Introduction

RNA plays essential roles in carrying genetic information that encodes protein sequences, as well as participating in catalysis and gene regulation as non-coding RNA (ncRNA)^1,2^. More recently, artificial RNAs have been designed and engineered as building blocks for nanostructure assembly and regulatory elements in cellular processes, which has enabled broad applications in nanomedicine and synthetic biology^3-6^. Riboswitches are a class of ncRNA elements that sense environmental changes and bind to ligands to regulate gene expression^7,8^. Their intrinsic regulatory functions facilitate design and engineering of novel riboswitches as biosensors, molecular reporters, and gene regulators in both prokaryotes and eukaryotes^9-13^.

T-box riboswitches bind to cognate tRNAs as ligands and sense their aminoacylation states in order to regulate transcription and translation of genes that control tRNA aminoacylation and amino acid levels in bacteria^14,15^. The majority of T-box RNAs utilize a specific codon, termed the Specifier Sequence, embedded in a complex Stem I helical domain that directs recognition of the cognate tRNA via complementary base pairing with the anticodon^15-18^. One class of translational T-box RNAs, termed ultrashort (US), replaces the normal Stem I with a much shorter helical element that contains the Specifier Sequence in the terminal loop; all of the T-box RNAs in this class regulate expression of *ileS* genes, encoding isoleucine-tRNA (tRNA^Ile^) synthetase, and respond to tRNA^Ile^. Long Stem I elements also contain a double T-loop structure in the distal end that stacks with the tRNA elbow^19-21^; this domain is missing in the US T-box RNAs. Most but not all T-box RNAs include a Stem II domain with a conserved S-turn that forms tertiary contacts with the Specifier Sequence to create a high-affinity binding pocket for tRNA^15-18,22^, a Stem IIA/B pseudoknot^16-18^, and a Stem III domain^16,20,23,24^. Transcriptional T-box RNAs contain a final region that forms competing terminator/antiterminator structures, whereas translational T-box RNAs contain competing sequestrator/antisequestrator (antiS) structures; in both cases, these elements determine downstream gene expression by controlling transcription attenuation or translational initiation, respectively. The Stem III domain stacks on the antiterminator or antisequestrator to facilitate sensing of tRNA aminoacylation^16,20,23,24^. Recent structural studies in transcriptional and translational T-box-tRNA complexes have provided detailed insights that may facilitate design and engineering of novel T-box-tRNA systems with enhanced regulatory functions^16,18-21^.

Advances in cryogenic-electron microscopy (cryo-EM) single particle analysis (SPA) have enabled determination of protein-free RNA structures at near-atomic resolution^25-27^. Here we use cryo-EM SPA to determine structure of the wild-type (WT) *Mycobacterium smegmatis* (MS) *ileS* T-box RNA in complex with its cognate tRNA^Ile^, and observe a very flexible antiS region with relative poor density that tolerates both 3’-OH and 2’,3’-cyclic phosphate (2’,3’-cP) at the 3’ end of tRNA, consistent with the *Mycobacterium tuberculosis* (Mtb) *ileS* T-box^16^. This finding facilitates our design to rigidify the antiS region by addition of one helical turn (11-base pair, 11-bp) to the acceptor arm of MS tRNA^Ile^ (acc.11-tRNA) and Stem III of MS *ileS* T-box RNA (S3.11-T-box) to increase base stacking in the antiS-tRNA region. Cryo-EM study demonstrated the formation of all of the T-box-tRNA complexes, and both the engineered T-box and tRNA increased the selectivity for 3’-OH over 2’,3’-cP at the tRNA’s 3’ end compared to WT T-box RNA and tRNA, as demonstrated by binding and *in vitro* translation results. In particular, S3.11-T-box RNA binds to WT tRNA with ∼6-fold higher binding affinity that resulted in precedent translation initiation compared to other T-box-tRNA complexes, indicating that Stem III stacking plays a critical role in tRNA recognition and translation regulation. These results demonstrate the potential of using cryo-EM to guide RNA engineering for novel regulatory modules with higher selectivity and sensitivity in synthetic biology.

## Results

### Cryo-EM reconstructions and models of WT and engineered MS *ileS* T-box-tRNA^Ile^ complexes

We first determined the cryo-EM structure of WT MS T-box in complex with its cognate uncharged tRNA. Although the complex is only 70 kDa, it was readily visible in the cryo-EM micrographs and two-dimensional (2D) averages (Extended Data Figure 1a-b). 3D reconstruction yielded the final cryo-EM structure at 6.32 Å global resolution (Figure 1a, Extended Data Figure 1c-e, Extended Data Table 1). Cryo-EM model and secondary structure were modified based on previous crystal structures of *ileS* T-box-tRNA^Ile^ complexes (Figure 1b)^16,18^. Local resolution map shows interrupted density of the antiS and Stem III domains indicative of high flexibility (Extended Data Figure 1e), which is in agreement with the low selectivity on tRNA 3’ end modification according to crystal structures of Mtb *ileS* T-box-tRNA^Ile^ complex^16^. Previous studies have suggested that insertion of one helical turn in tRNA acceptor arm could fully restore glyQS T-box antitermination activity^28,29^. We rigidified the antiS and Stem III domains by insertion of an additional 11-bp helical turn in Stem III of MS T-box (S3.11-T-box) and acceptor arm of MS tRNA^Ile^ (acc.11-tRNA) in order to enhance selectivity and sensitivity of tRNA recognition. We subsequently determined cryo-EM structures of three complexes, including T-box-acc.11-tRNA at 7.50 Å resolution, S3.11-T-box with WT tRNA and acc.11-tRNA at 9.63 Å and 6.85 Å resolution, respectively (Figure 1c-h, Extended Data Figure 1f-t). Because antiS and Stem III domains were generally not well resolved for all T-box-tRNA complexes after 3D refinement in cryoSPARC^30^ (Extended Data Figure 2), we performed one round of 3D classification in Relion^31^ that focus on antiS and Stem III domains and yielded final cryo-EM maps with improved density in the antiS and Stem III region (Figure 1, Extended Data Figure 2c). Better resolved local maps of antiS and Stem III in T-boxes complexed with acc.11-tRNA reveal that elongated acceptor arm helped to rigidify the antiS and Stem III domains (Figure 1c, 1g).

**Figure 1.**
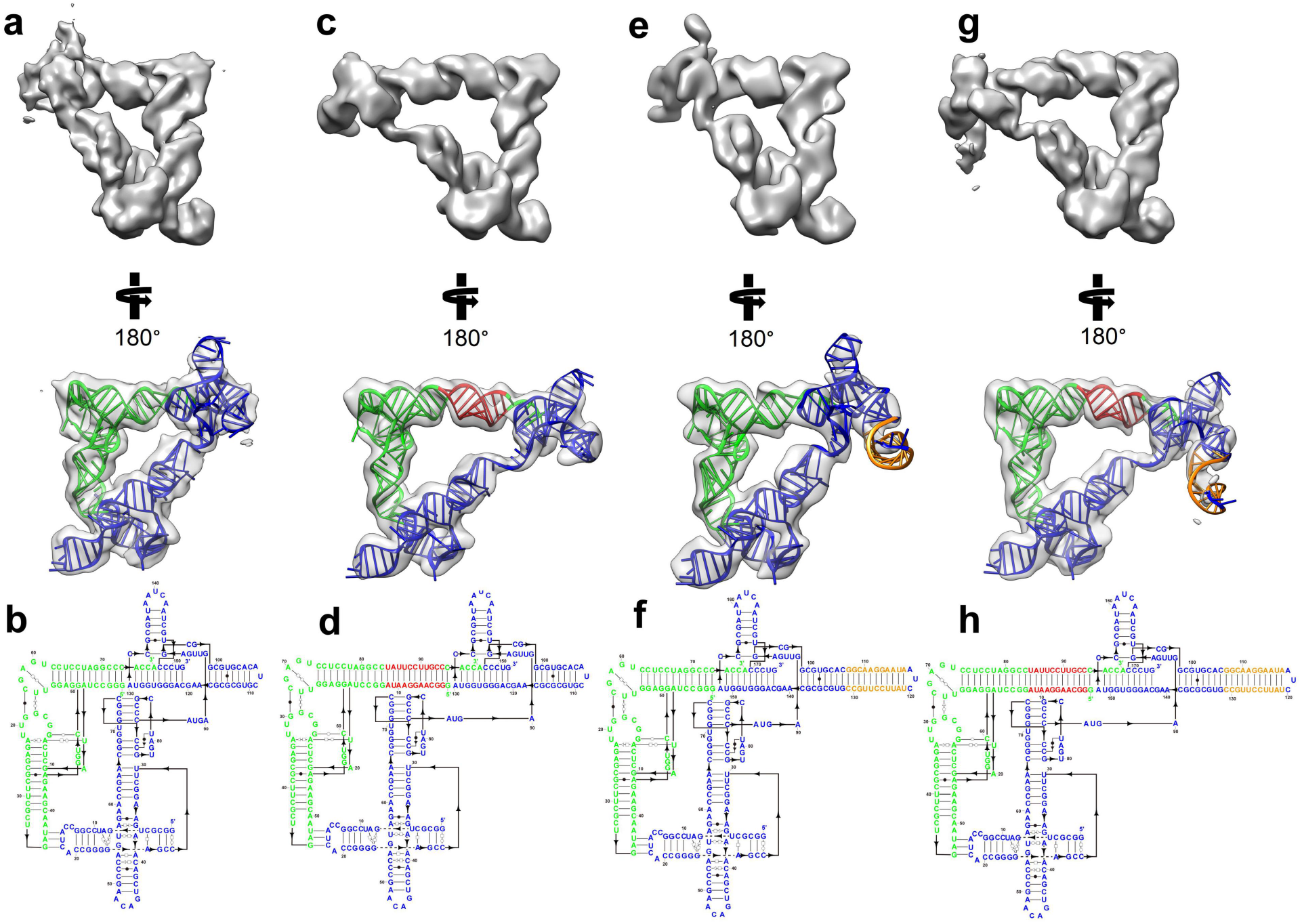
Cryo-EM structures and models of WT and engineered MS *ileS* T-box-tRNA^Ile^ complexes. Cryo-EM maps and models, and secondary structures of WT T-box-tRNA (a-b), T-box-acc.11-tRNA (c-d), S3.11-T-box-tRNA (e-f), and S3.11-T-box-acc.11-tRNA (g-h). Green region indicates WT tRNA, blue region indicates WT T-box, red region indicates inserted stem in acc.11-tRNA, orange region indicates inserted stem in S3.11-T-box.

**Figure 2.**
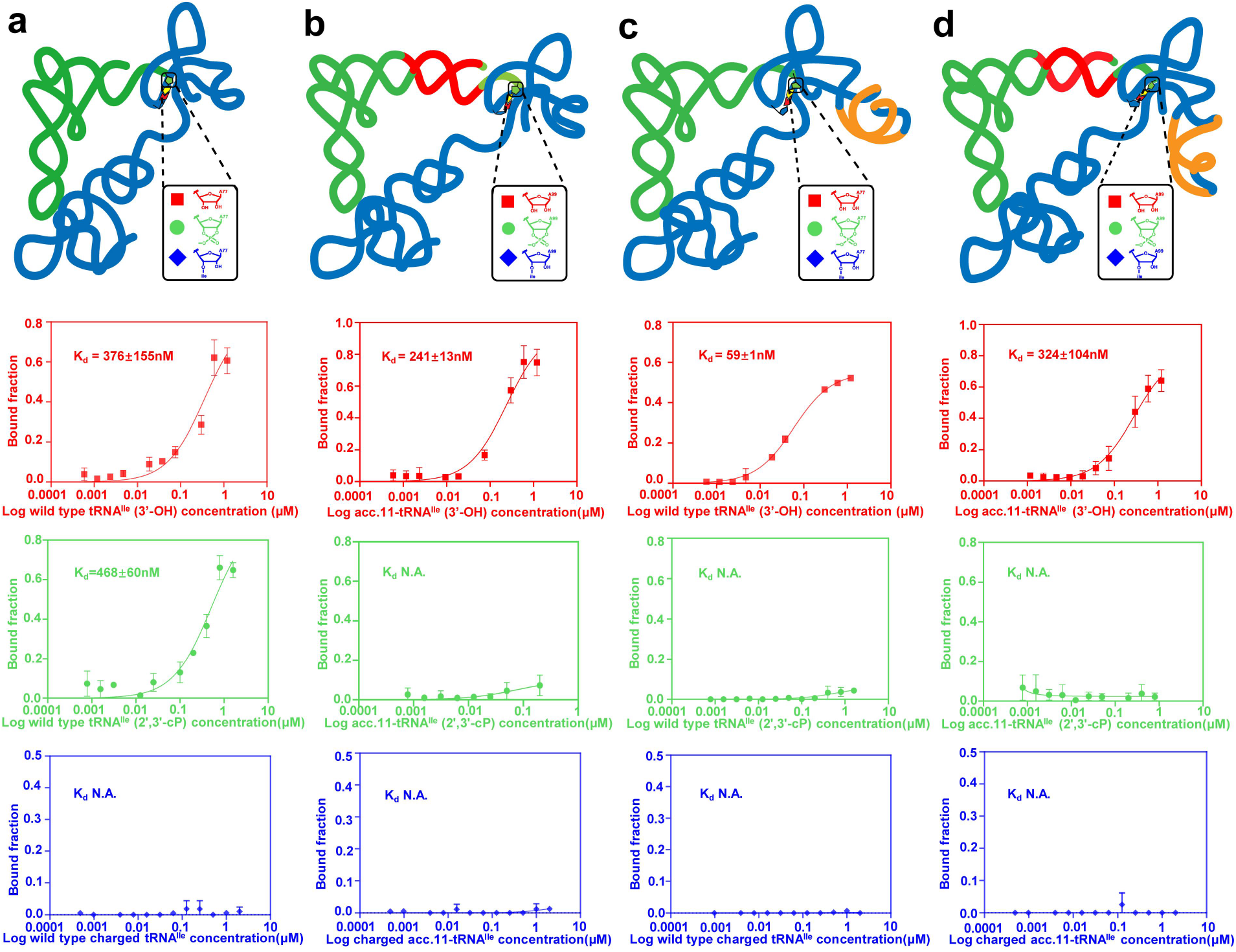
Binding affinity of WT and engineered MS *ileS* T-box-tRNA^Ile^ complexes with different 3’ end modifications on tRNAs. (a) Binding affinity of WT T-box and tRNA with different 3’ modifications. (b) Binding affinity of WT T-box and acc.11-tRNA with different 3’ modifications. (c) Binding affinity of S3.11-T-box and WT tRNA with different 3’ modifications. (d) Binding affinity of S3.11-T-box and acc.11-tRNA with different 3’ modifications. 3’ modifications include 3’-OH (red square), 2’,3’-cP (green circle) and 3’-aminoacylation (blue diamond). Green cartoon region indicates WT tRNA, blue cartoon region indicates WT T-box red cartoon region indicates inserted stem in acc.11-tRNA, orange cartoon region indicates inserted stem in S3.11-T-box.

### Engineered tRNA acceptor arm and T-box Stem III increase T-box selectivity on 3’ end modifications of tRNAs

Cyclic phosphate modification on tRNA’s 3’ end is of similar size as the smallest amino acid glycine, which is frequently used to evaluate steric hindrance of the T-box regions that bind to the tRNA’s 3’ end^16,20,24^. Previous studies on transcriptional *glyQS* T-box suggested a steric occlusion mechanism that discriminate 2’,3’-cP from 3’-OH^20,24^. In contrast, translational Mtb *ileS* T-box antiS domain has been found to tolerate both 3’-OH and 2’,3’-cP^16^. In order to assess if engineering on T-box and tRNA has impact on steric hindrance of tRNA’s 3’ end when binding to MS T-box, we prepared both WT and acc.11-tRNA with different 3’ end modifications of 3’-OH, 2’,3’-cP and 3’-aminoacylation of isoleucine (3’-ile).

We first evaluated bindings of WT T-box with tRNAs of different 3’ end modifications and observed complex formations for WT T-box with tRNAs of 3’-OH and 2’,3’-cP, but not of 3’-ile (Figure 2a, Extended Data Figure 3a-c)^16^. Next, WT T-box was found to have higher selectivity on 3’ end modifications of acc.11-tRNA that discriminated 2’,3’-cP and 3’-ile from 3’-OH, indicative of more compact antiS binding region with higher steric hindrance in the presence of additional 11-bp in the tRNA acceptor arm (Figure 2b, Extended Data Figure 3d-f). Adding 11-bp in the T-box Stem III had similar enhancement in selectivity of tRNA 3’ end modifications that S3.11-T-box only bound to uncharged WT and acc.11-tRNA (Figure 2c-d, Extended Data Figure 3g-l). Finally, all T-boxes bound to tRNAs with similar binding affinity, except for S3.11-T-box-tRNA complex, which was formed with ∼6-fold higher affinity (Figure 2c). Although neither the sequence nor the length of Stem III is conserved^32^, recent studies suggested that Stem III could stack on and stabilize antiterminator/antiS domain^16,20^, which may explain the enhanced tRNA binding affinity by the additional 11-bp stacking on the antiS domain.

**Figure 3.**
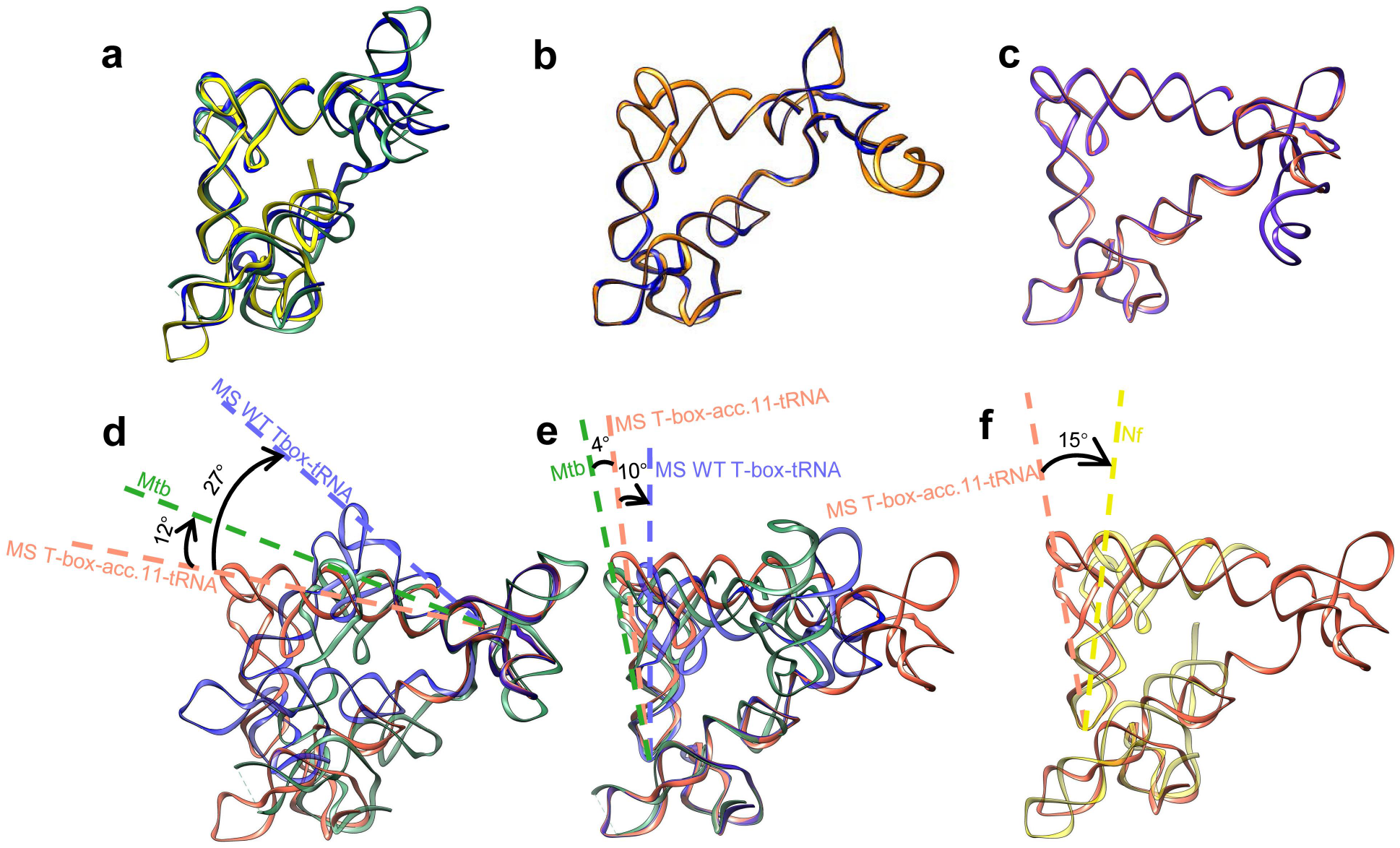
Superposition of cryo-EM structures of WT and engineered MS *ileS* T-box-tRNA^Ile^ complexes with crystal structures of different *ileS* T-box-tRNA complexes, aligned at Specifier loops or antiS regions. (a) Superposition of WT MS (blue), Mtb (green) and Nf (yellow) *ileS* T-box-tRNA complex. (b) Superposition of MS S3.11-T-box-tRNA (orange) and T-box-tRNA (blue). (c) Superposition of MS S3.11-Tbox-acc.11-tRNA (purple) and T-box-acc.11-tRNA (salmon). (d-f) The MS T-box-acc.11-tRNA complex (salmon) is superimposed with MS T-box-tRNA complex (blue) and Mtb T-box-tRNA complex (green) aligned at antiS regions (d) and Specifier loop regions (e), with Nf T-box-tRNA complex (yellow) aligned at Specifier loop regions (f).

### acc.11-tRNA adopts deviated trajectories on both acceptor and anticodon arms compared to WT tRNA in T-box-tRNA complex

Superposition of our cryo-EM model of WT MS *ileS* T-box-tRNA and previous crystal structures of Mtb and *Nocardia farcinica* (Nf) *ileS* T-box-tRNA complexes shows overall agreement on the global architecture (Figure 3a). MS WT T-box and S3.11-T-box also adopt almost identical structures when complexed to the same tRNA, except for the elongated helical turn in Stem III stacks on the antiS of S3.11-T-box (Figure 3b-c). The acc.11-tRNA forms a sharper angle at the elbow region to maintain interactions with both antiS and Specifier loop of the WT T-box, resulting in deviations of the acceptor arm up to 27° from WT tRNAs when aligned to antiS regions (Figure 3d), and deviations of the anticodon arm up to 15° from WT tRNAs when aligned to Specifier loops (Figure 3e-f). The acc.11-tRNA complexed with S3.11-T-box adopts similar deviations of the acceptor and anticodon arms compared to WT tRNAs (Extended Data Figure 4).

**Figure 4.**
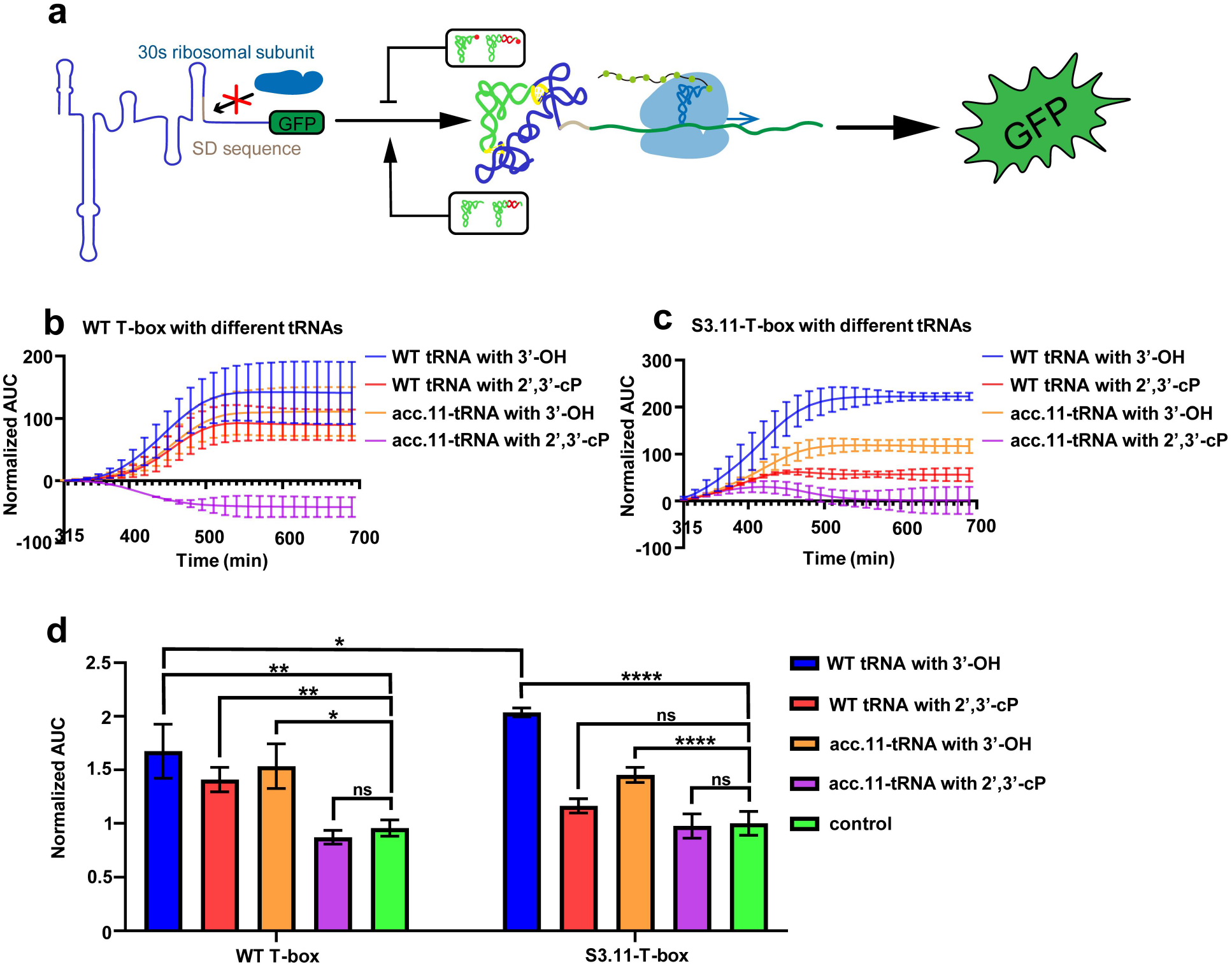
Translation reporter assay of WT and engineered *ileS* T-box-tRNA^Ile^ complexes. (a) Schematic illustration of the *in vitro* translation regulation. (b-c) Normalized integrations of GFP expression curves of WT T-box (b) and S3.11-T-box (c) in the presence of different tRNAs. (d) End-point quantifications of *in vitro* translation regulations of WT T-box and S3.11-T-box in the presence of different tRNAs including uncharged WT tRNA (blue), WT tRNA with 2’,3’-cP (red), uncharged acc.11-tRNA (orange), acc.11-tRNA with 2’,3’-cP (purple), and no tRNA (green) as negative control. All experiments were performed in triplicate or quadruplicate and data are represented as mean values ± SD. ns P>0.05; * P≤0.05; ** P≤0.01; *** P≤0.001; **** P≤0.0001 (two-tailed T test).

### Engineered MS *ileS* T-box and tRNA^Ile^ present improved selectivity and sensitivity in downstream translational regulations

Formation of MS *ileS* T-box-tRNA^Ile^ complex regulates downstream gene expression^15^. We designed *in vitro* reporter assay to evaluate the impact of engineered MS T-box and tRNA on green fluorescent protein (GFP) expression. WT *ileS* T-box and S3.11-T-box sequences were individually inserted at the 5’ end of GFP sequence, tRNA^Ile^ with 3’-OH or 2’,3’-cP modifications were then added to the *in vitro* translation system and the expressed GFP signal was monitored (Figure 4a). The resulting normalized GFP signals were quantified by integration as translation proceeded to evaluate impact of different tRNAs on translation initiations (Figure 4b-c, Extended Data Figure 5, see Materials and Methods for details). WT T-box could bind to both WT tRNAs with 3’-OH and 2’,3’-cP to accelerate the translation initiations, whereas the remaining combinations of engineered T-box-tRNA complexes required uncharged tRNAs to accelerate translation initiations (Figure 4b-d), demonstrating the increased selectivity of engineered riboswitch systems. Intriguingly, S3.11-T-box significantly accelerated translation initiation in the presence of uncharged WT tRNA compared to WT T-box, indicating that higher binding affinity may correspond to increased sensitivity of tRNA recognition.

In conclusion, we have designed artificial translational MS *ileS* T-box and tRNA^Ile^ with enhanced selectivity and sensitivity to tRNA recognition and downstream gene expression regulation. This study demonstrates the potential of cryo-EM to guide design of improved regulatory RNA modules for therapeutics, nanotechnology and synthetic biology.

## Discussion

T-boxes are mostly found in the 5’ untranslational regions of the aminoacyl-tRNA synthetase coding genes to bind to cognate tRNAs and regulate downstream transcriptions and translations^14,15^. Recent structural studies of both transcriptional and translational T-boxes complexed with cognate tRNAs revealed complete molecular mechanisms of tripartite and bipartite tRNA recognitions, and discriminations of tRNA 3’ end modifications^16,18-21^. Unlike the transcriptional *glyQ* T-boxes that occlude charged tRNA with modification as small as 2’,3’-cP by steric hindrance^20^, the translational *ileS* T-box could bind to tRNA with 2’,3’-cP and undergo deconformation of the antiS to form sequester conformation^16^.

Recent advances in RNA cryo-EM structure determination allow us to resolve cryo-EM structures of WT and engineered MS *ileS* T-box-tRNA^Ile^ complexes with 11-bp insertions in the tRNA acceptor arm and T-box Stem III domains (Figure 1)^27,33^. The insertions rigidify the antiS-tRNA complex structure and change the tRNA recognition mechanism from antiS deconformation to steric hindrance in order to provide more selective recognitions of uncharged tRNAs, as demonstrated by discrimination of 2’,3’-cP from 3’-OH on the tRNA’s 3’ end in the binding experiment (Figure 2). Due to the enhanced selectivity, engineered T-box-tRNA complexes present riboswitch systems that regulate downstream translation more precisely than the WT system, as shown by the *in vitro* translation reporter assay (Figure 4). Additional stacking of engineered Stem III on antiS is found to enhance binding of WT uncharged tRNA that results in increased sensitivity in tRNA recognition of S3.11-T-box (Figure 4d). Our study reveals the capability of cryo-EM SPA to facilitate the design of improved RNA regulatory modules potentially useful for broad RNA applications.

## Materials and Methods

### RNA preparations

RNA was prepared as previously described^17^. Briefly, the DNA template was amplified from the pUC19-T7-template-HDV. The DNA templates of all tRNAs with 2’,3’-cP modifications on the 3’ end were linearized plasmids containing HDV ribozyme sequence in order to generate homogeneous 2’,3’-cP modifications on the RNAs’ 3’ ends^27,34^. The DNA templates of all RNAs with 3’-OH were amplified by PCR with reverse primers containing two 2’-O-Methyl modified bases at the 5’ end to yield homogeneous 3’-OH on the RNAs’ 3’ ends^35^. The RNA was prepared by *in vitro* transcription in a reaction containing 0.2 μM DNA template, 40 mM Tris-HCl, pH=7.9, 20 mM MgCl^2^, 2 mM spermidine, 0.01% TritonX-100, 10 mM DTT, 4 mM NTPs, and 0.8 μg/μL T7 RNA polymerase. The transcription reaction was incubated at 37 °C for 3-4 h. The RNA was then centrifuged at 13,000 rpm for 5 min and the supernatant was mixed with loading buffer containing 6M urea, 1×Tris/Borate/EDTA buffer, 0.1% xylene cyanol, and 0.1% bromophenol blue and loaded on an 8% 29:1 acrylamide:bis, 8 M urea polyacrylamide gel. The gel electrophoresis was performed at 300V for 2-3 h, then visualized briefly with a 254-nm UV lamp, held far from the gel to minimize RNA damage^36^. RNA was eluted from the gel overnight in 30 mM NaOAc pH=5.2, 0.1 mM EDTA at 4 °C, then concentrated with Amicon® Ultra-15 3K Centrifugal Filter Devices (Milipore) to a total volume of 500 μL and precipitated with isopropanol.

### Aminoacylated tRNA^Ile^ preparation

Charged tRNA^Ile^ was prepared by an *in vitro* method using eFx (enhanced flexizyme) to catalyze the acylation of L-isoleucine 4-chlorobenzylthioester (ile-CBT) on the 3’ end of tRNA^Ile37^. The acylation reaction was performed as follow: 40μM WT tRNA^Ile^ or acc.11-tRNA^Ile^ (3’-OH) in 200 mM HEPES-KOH pH=7.5 was heated at 95 °C for 3 min and cooled to room temperature over 5 min. 1μL MgCl^2^ (3 M) and 0.5 μL eFx (200 μM) were added. The mixture was incubated at room temperature for 5 min, then 1μL of 25mM ile-CBT (in DMSO) was added to initiate the reaction and proceeded on ice for 6 h. Finally, 15 μL of 0.6M NaOAc (pH=5.0) was added to terminate the reaction and the mixture was mixed with loading buffer containing 100 mM NaOAc pH=4.7, 0.05% xylene cyanol and 6M urea. Electrophoresis was performed in running buffer of 100mM NaOAc (pH=4.7) on a 12% polyacrylamide gel (29:1 acrylamide:bis, 8 M urea, 50mM NaOAc, pH=4.7). The purification was completed by gel extraction, elution and precipitation as mentioned in RNA preparations above.

### Electrophoresis mobility shift assay (EMSA)

Purified RNA concentration was estimated using the A280/260 value on the NanoDrop One Microvolume UV-Vis Spectrophotometer (Thermo Scientific). All tRNAs were heated to 85 °C for 5 min, cooled to room temperature to allow refolding in binding buffer (20mM Tris-HCl pH=7.9, 20mM NaCl, 10mM MgCl^2^, 0.1mM EDTA). Then, tRNAs were aliquoted by 1:2 serial dilutions and T-box (100 nM final concentration) was added to each aliquot to a final volume of 5 μL. All tRNAs with 3’-OH were diluted from 1.2 μM. All tRNAs with 2’,3’-cP were diluted from 1.6 μM. Charged tRNAs were diluted from 2 μM. Reaction mixtures were heated to 65 °C for 5 min and cooled to room temperature. Reactions were instantly placed on ice and 5 μL native loading buffer (10mM Tris-HCl pH=7.4, 100mM KCl, 10mM MgCl^2^ 0.05% xylene cyanol, 0.05% bromophenol blue) was added before loading onto a native 6% polyacrylamide gel (29:1 acrylamide:bis, 50 mM Tris-HCl pH=7.9, 100mM KCl, 10mM MgCl^2^). Electrophoresis was performed at 4 °C at 120 V for 40 min in 1× THE buffer (33 mM Tris base, 300 mM HEPES, 0.1 mM EDTA pH=8.0). Gels were stained by SYBR™ Gold Nucleic Acid Gel Stain (Invitrogen), imaged by ChemDoc XRS+ (BioRad), quantified using ImageJ. The experiments of WT T-box-WT tRNA with 3’-OH, S3.11-T-box-WT tRNA with 3’-OH or 2’,3’-cP, WT or S3.11-T-box-charged tRNA were repeated with two independent RNA isolations and the other reactions were repeated with three independent RNA isolations. Dissociation constants (Kd) were calculated by one site-specific binding of nonlinear regression analysis using GraphPad Prism9.0.0 software.

### Construction of vectors for *in vitro* expression assays

All T-box and GFP fusions were generated from plasmid pUC19 carrying the entire WT and S3.11-T-box (containing entire sequences upstream initiation codon including SD sequences) and superfold GFP reporter gene.

The required insertion (T7 promoter-Tbox-GFP-T7 terminator) for homologous recombination was amplified by overlap extension PCR from the plasmids described above using appropriate oligonucleotide primers. For GFP expression control, only superfold GFP gene was inserted between T7 promoter and T7 terminator. For T-box *in vitro* translations, the entire WT T-box or S3.11-T-box was placed immediately upstream of the superfold GFP reporter gene through overlap extension PCR.

### *In vitro* translation reporter assay

Fluorescence intensity measurements at excitation of 479 nm and emission of 520 nm were performed using a BioTek Cytation 5 Cell Imaging Multi-Mode Reader in a 96-well plate. Each reaction system contained 10.3 μL reaction mix and 4.4 μL *E*.*coli* cell extract from the cell-free protein expression kit (GZL #CF-EC-1000B), and DNA template with a final concentration of 33 nM. The mixture was stored on ice and adjusted to a total volume of 40 μL using RNase free water and the corresponding tRNA 12.5 μM final concentration. Additional GFP expression control reactions in the presence or absence of corresponding tRNAs were prepared to eliminate background effects from different tRNAs on GFP expression. The final reaction was then added into a 96 well plate and recorded at 15 min intervals with 10 s linear shaking before each read for a total of 47 reads at 37 °C. The plotted GFP intensity over time was normalized among different reactions, and the area under curve (AUC) value was integrated for each reaction starting from 315 min when the baseline became steady (Extended Data Figure 5a-f, Extended Data Figure 6a-b). The T-box-GFP AUC curves over time were then subtracted by GFP curves in the presence of corresponding tRNAs, followed by subtractions of each resulting normalized T-box-GFP curve with tRNA by the normalized T-box-GPF alone curve (Extended Data Figure 5g-m, Extended Data Figure 6d). The final normalized AUC curves of T-box-GFP with corresponding tRNAs over time represent the effect of each tRNA on T-box-GFP translation initiation. The quantitative histograms were plotted using the end points in the initially normalized T-box-GFP curves in the presence or absence of tRNAs divided by the end point in the corresponding GFP curves. The result in the absence of tRNAs was normalized to 1 as the control (Extended Data Figure 6c). All curves were plotted in Graphpad Prism 9.0.0, the values were the means ± s.d. of n=3 or 4 biologically independent experiments.

### Cryo-EM sample preparation

A total of 3 μL of MS *ileS* T-box and tRNA^Ile^ sample (25 μM T-box to 25 μM tRNA^Ile^ mixture treated as described in EMSA was applied onto glow-discharged (40 s) 200-mesh R2/2 Quantifoil Cu grids. The grids were blotted for 1.5 s in 100% humidity with no blotting offset and rapidly frozen in liquid ethane using a Vitrobot Mark IV (Thermo Fisher).

### Cryo-EM single particle data acquisition and data processing

The abovementioned frozen grids were loaded in Titan Krios (Thermo Fisher) operated at 300 kV, condenser lens aperture 50 μm, spot size 7, parallel beam with illumine area of 1.05 μm in diameter. Microscope magnification was at 130,000× (corresponding to a calibrated sampling of 1.06 Å per physical pixel). Movie stacks were collected automatically using EPU software on a K2 Bioquantum direct electron device equipped with an energy filter operated at 20 eV (Gatan), operating in counting mode at a recording rate of 5 raw frames per second and a total exposure time of 5 seconds, yielding 25 frames per stack, and a total dose of 49 e^−^/Å^2^. For WT MS *ileS* T-box and WT tRNA^Ile^ complex dataset, two datasets were collected and a total of 5,792 movie stacks were collected with defocus values ranging from -2.46 to -3.26 μm. These movie stacks were motion corrected using Motioncor2^38^. After CTF correction by CTFFIND4^39^, then subjected to EMAN2.2 for neural network particle picking^40^. A total of 3,219,341 particles were extracted in Relion3^31^. After two independent rounds of 2D classifications of two datasets, the best classes were selected by visual examination. The smaller dataset was subjected to cryoSPARC3.2.0 to build initial models and select the best class for non-uniform refinement. Then, a total of 830,281 particles were subjected to cryoSPARC3.2.0 to build the initial model, but didn’t obtain cryo-EM map with RNA features. Therefore, the total particles were subjected to heterogeneous refinement in cryoSPARC3.2.0 using the above non-uniform refinement result as an extra input initial model. The major class showing RNA features, including 152,679 particles, were subjected to cryoSPARC homogeneous refinement and Relion focus classification on the antiS region. Finally, the best class was subjected to homogeneous refinement in cryoSPARC3.2.0 to yield the final map. A sharpening B-factor of -377.6 Å^2^ was applied to the resulting cryo-EM map to yield the final sharpened map at 6.32 Å resolution estimated by the 0.143 criterion of FSC curve. The other three complex dataset, the data processing was similar and detailed parameters were showed in Extended Data Table1. The appropriate low-pass filters were applied to the final 3D map displayed in UCSF Chimera^41^.

### Cryo-EM model building and refinement

The initial MS *ileS* Tbox-tRNA^Ile^ complex model were built with auto-DRRAFTER^5^, then manually adjusted and rebuilt with Coot as needed^42^. The models were refined with Phenix.real_space_refine^43^, yielding model–map correlation coefficient (CCmask) of 0.75 (WT T-box-tRNA), 0.75 (WT Tbox-acc.11-tRNA), 0.59 (S3.11-T-box-WT tRNA) and 0.80 (S3.11-T-box-acc.11-tRNA). The final model was validated by MolProbity^44^. Secondary structure diagrams were prepared with RiboDraw aided by manual adjustment. (https://github.com/ribokit/RiboDraw).

## Acknowledgements

Cryo-EM data were collected at SKLB West China Cryo-EM Center and SLAC-Stanford Cryo-EM Center, processed on Duyu High Performance Computing Center in Sichuan University. This work was supported by Ministry of Science and Technology of China (2022YFC2303700 and 2021YFA1301900 to Z.S.), the National Natural Science Foundation of China (32222040 and 32070049 to Z.S.), and Sichuan University start-up funding 20822041D4057 to Z.S., A.M.W. was supported by Army Research Office W911NF-16-1-0372 and RNA modeling computation was supported by National Institutes of Health R35 GM122575.

## Author Contributions

Z.S. and T.M.H designed experiments, X.J., C.Z. and J.K.F. prepared RNAs, X.J, Y.Y., C.Z. and J.K.F. performed biochemical and functional experiments, Z.S. collected cryo-EM data, X.J, B.L. and K.L. processed Cryo-EM data, Z.S., X.J., A.M.W. and B.L. built and refined atomic models. All authors contributed to preparation of the manuscript.

## Data availability

The cryo-EM map and associated atomic coordinate model of the WT and S3.11-T-box in complex with WT and acc.11-tRNA^Ile^ complexes have been deposited in the wwPDB OneDep System under EMD accession code EMD-35513, EMD-35515, EMD-35517, EMD-35517, and PDB ID code 8IKK, 8IKI, 8IKN, 8IKO. The authors declare no competing financial interests.

