## Supplementary information for "Cryo-EM-guided engineering of T-box-tRNA modules with enhanced selectivity and sensitivity in translational regulation"

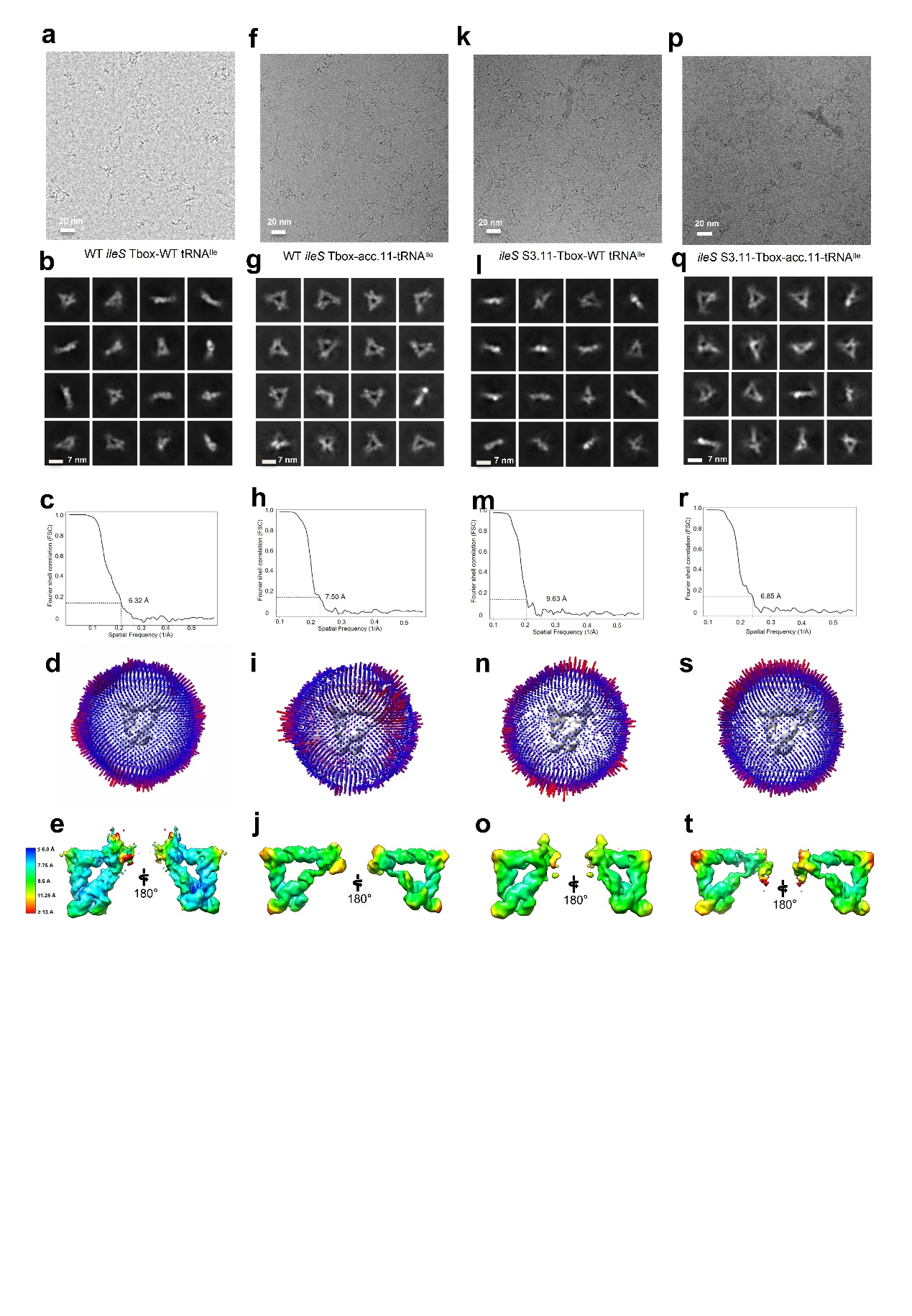

**Extended Data Figure 1. Cryo-EM SPA results of WT MS *ileS* T-box and S3.11-T-box complexed with WT tRNA^Ile^ and acc.11-tRNA^Ile^.** Cryo-EM micrographs, 2D class averages, FSC curves, angular distribution and local resolution maps are presented for WT T-box with tRNA complex (a-e), WT T-box with acc.11-tRNA (f-j), S3.11-T-box with WT tRNA complex (k-o) and S3.11-T-box with acc.11-tRNA complex (p-t).

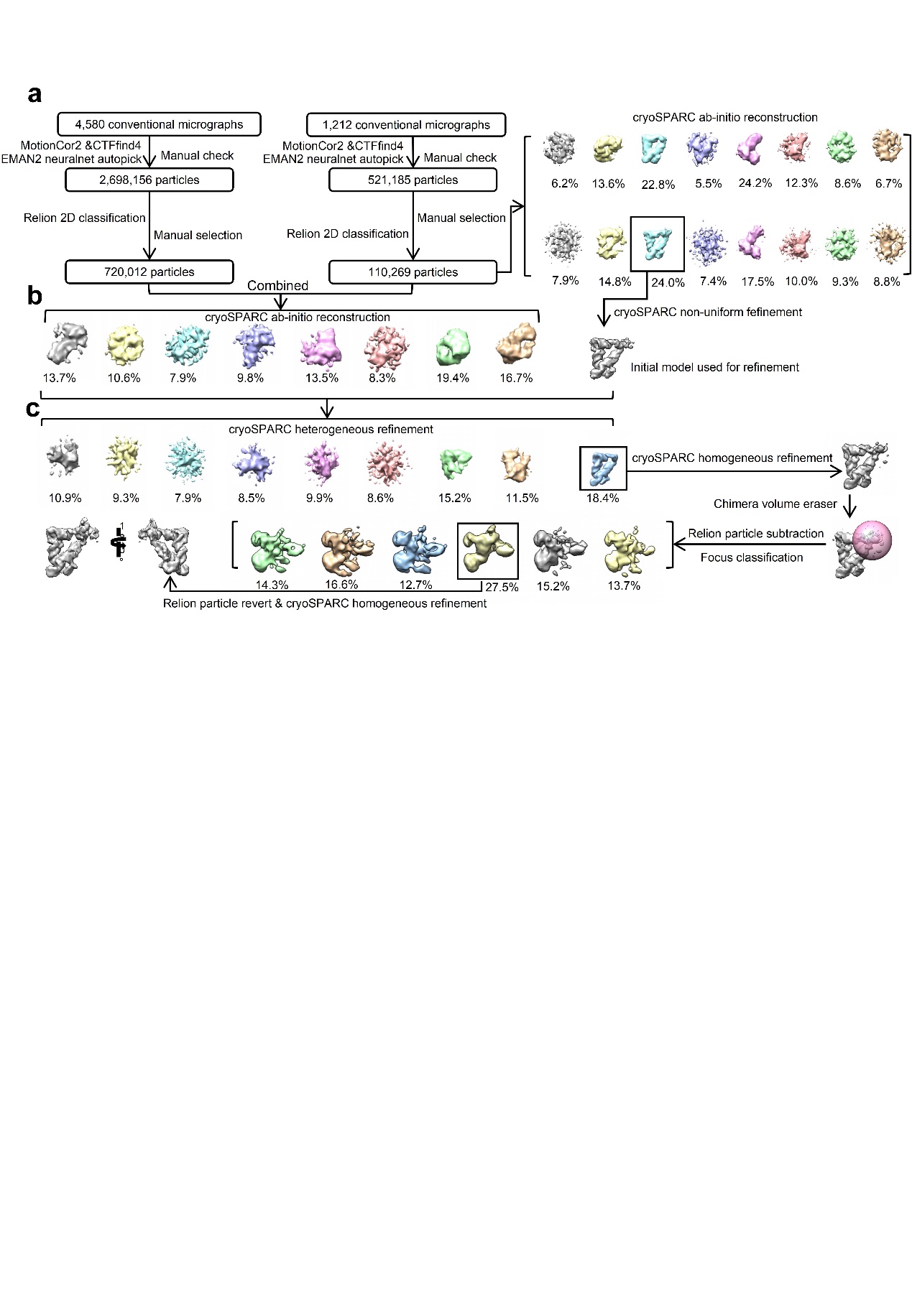

**Extended Data Figure 2. Representative Cryo-EM SPA workflow of WT T-box-tRNA^Ile^ complex.** (a) Two data sets were collected for WT T-box-tRNA^Ile^ complex. After 2D classification in Relion, the resulting particles from the smaller data set was subjected to cryoSPARC for ab-initio reconstruction to yield a major class (black box) with RNA features. (b) 2D classification results from two data sets were combined for cryoSPARC ab-initio reconstruction, which did not result in cryo-EM maps with RNA features. The cryoSPARC non-uniform refinement result from the smaller data set was then included as the input initial model for cryoSPARC heterogeneous refinement of the combined two data sets. (c) The resulting major class was subjected to cryoSPARC homogeneous refinement and Relion focus classification on the antiS region, which generated a major class (black box) for cryoSPARC homogeneous refinement to yield the final map.

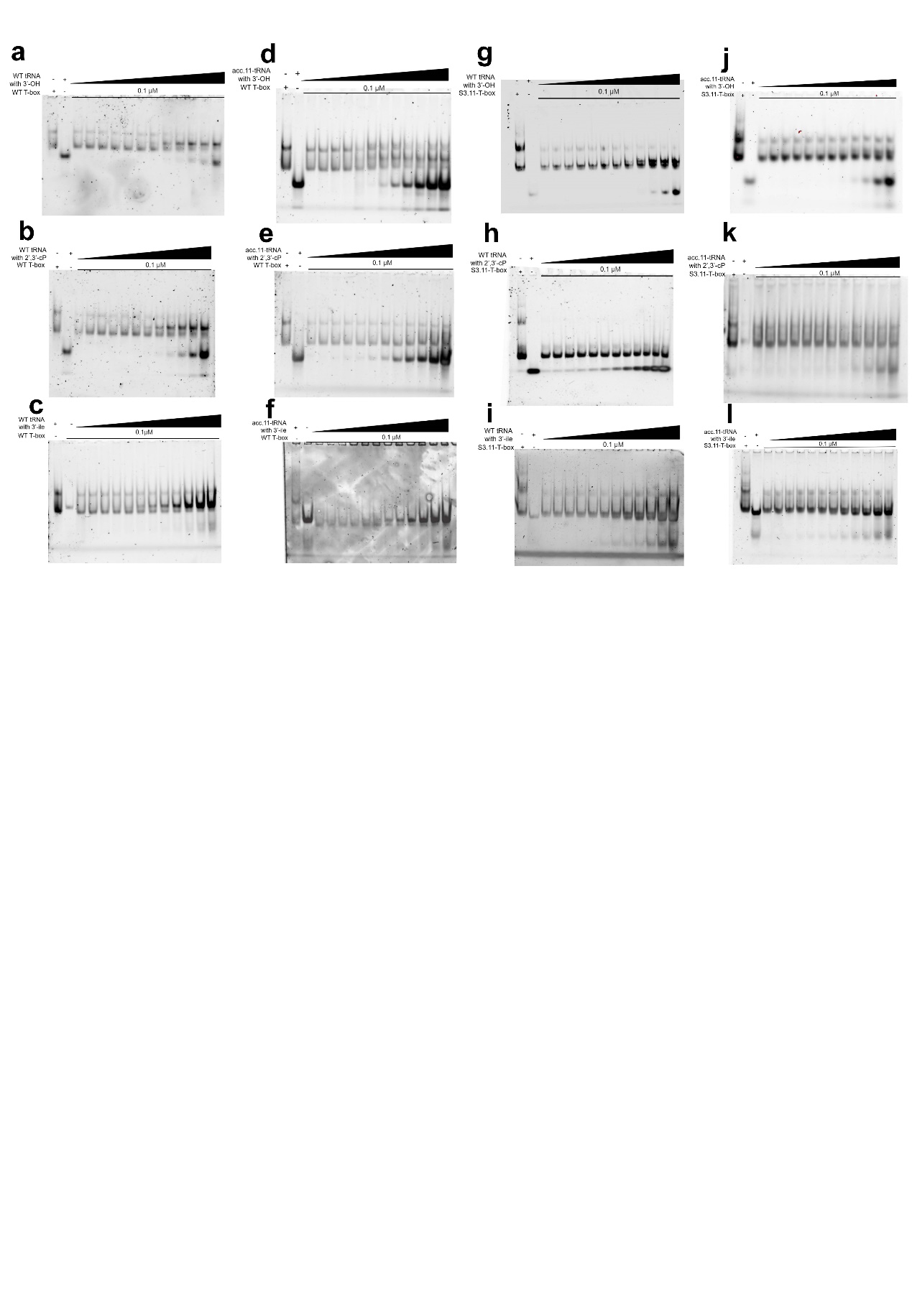

**Extended Data Figure 3. Representative EMSA analyses of WT and S3.11-T-box complexed with WT tRNA^Ile^ and acc.11-tRNA with different 3’ end modifications.** Complex formations of WT T-box and WT tRNA with 3’-OH (a), 2',3'-cP (b) and 3’-aminoacylation (c), and acc.11-tRNA with 3’-OH (d), 2',3'-cP (e) and 3’-aminoacylation (f) were examined by EMSA. Complex formations of S3.11-T-box and WT tRNA with 3’-OH (g), 2',3'-cP (h) and 3’-aminoacylation (i), and acc.11-tRNA with 3’-OH (j), 2',3'-cP (k) and 3’-aminoacylation (l) were examined by EMSA.

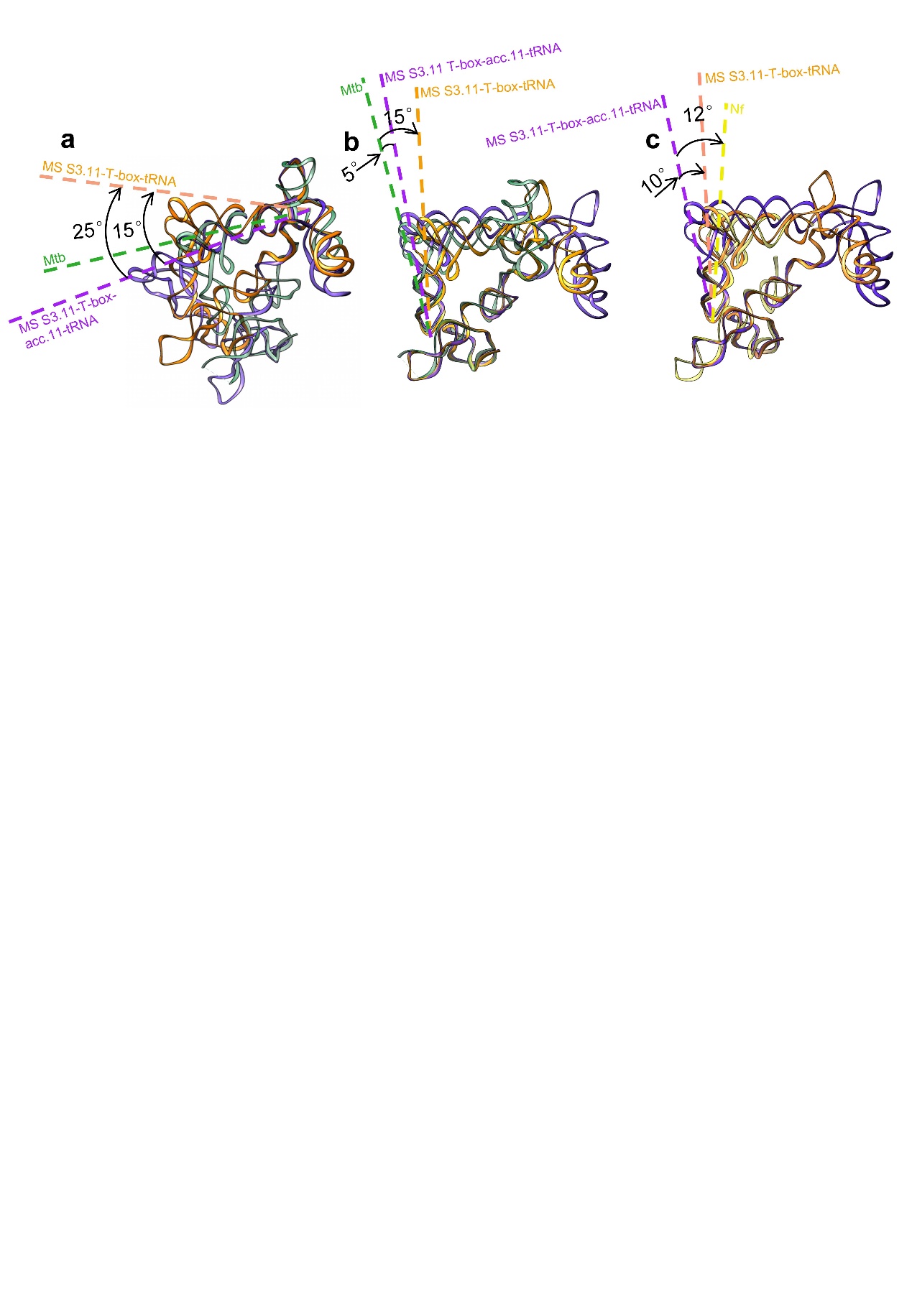

**Extended Data Figure 4. Superposition of cryo-EM structures of MS S3.11-T-box-tRNA complexes with crystal structures of different *ileS* T-box-tRNA complexes, aligned at specifier loops or antisequestrator regions.** The MS S3.11-T-box-acc.11-tRNA complex (purple) is superimposed with MS S3.11-T-box-tRNA complex (orange) and Mtb T-box-tRNA complex (green) aligned at antiS regions (a) and specifier loop regions (b), with Nf T-box-tRNA complex (yellow) aligned at specifier loop regions (c).

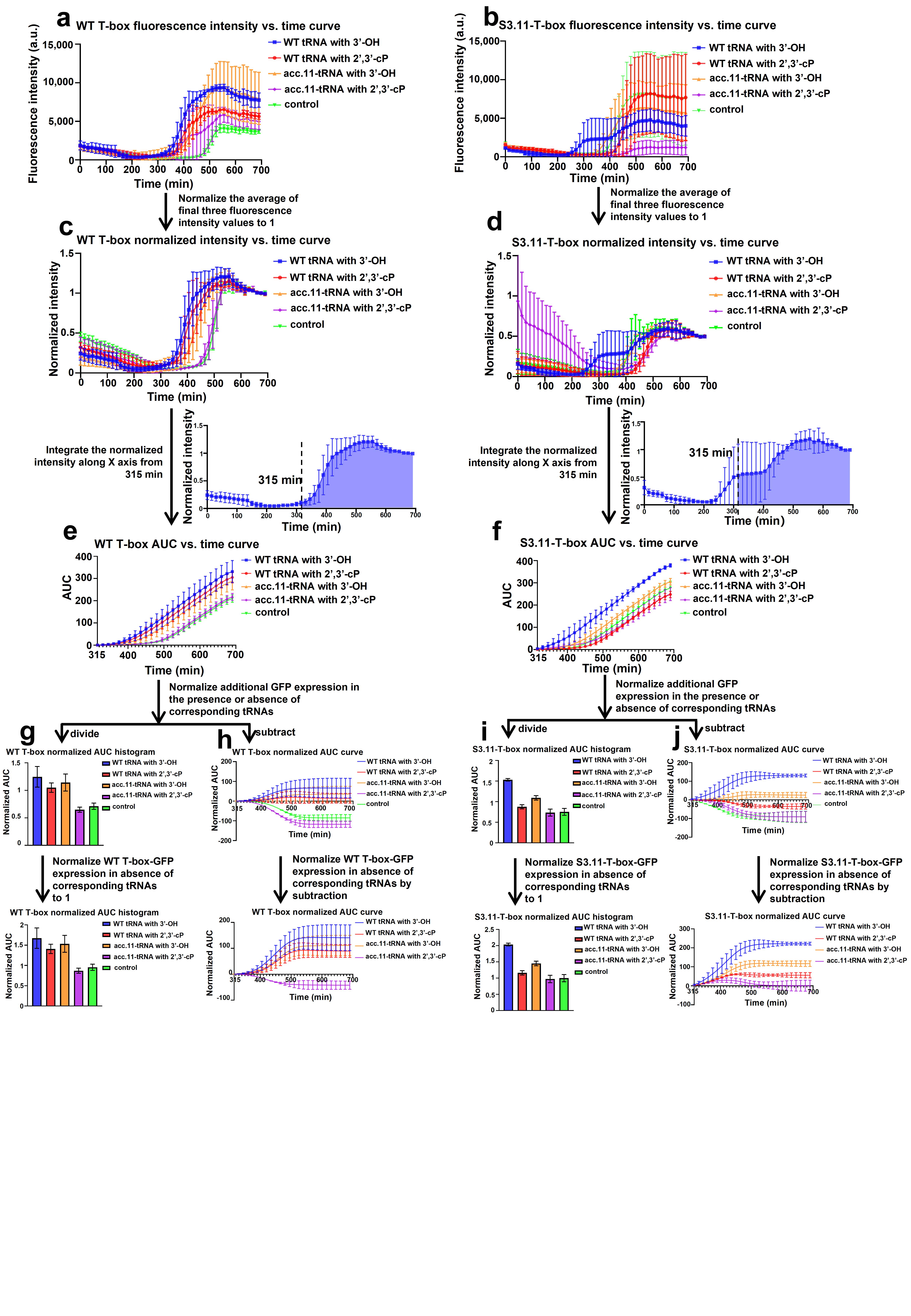

**Extended Data Figure 5. Representative in vitro expression analysis workflow of WT complexed with WT tRNA^Ile^ and acc.11-tRNA with different 3’ end modifications.** The fluorescence intensity of WT T-box with different tRNAs and additional GFP expression control reactions were collected and plotted over time (a). Then the mean value of last three intensities were normalized to 1 (b). The AUC curves were calculated based on the area under curves (AUC) along the X-axis of normalized intensity curves from 315 min (c). Normalized AUC curves were plotted using the subtractions of WT T-box AUC curves and the mean value of the additional GFP expression reactions (d). The mean value of tRNA free translations were then subtracted from subtraction values of tRNA-treated translations to obtain normalized subtraction curves (e). The histograms were plotted based on the AUCs of the normalized intensity vs. time curves of WT T-box translations from 315 min (normalized through dividing the AUC values of the initial curves by additional GFP expression control reactions). Thereinto, the mean value of tRNA free reaction was normalized to 1 (f).

Table S1. Cryo-EM data collection, processing, and model refinement statistics of WT and engineered MS *ileS* T-box-tRNA^Ile^ complexes.

| **Cryo-EM data collection and processing** | WT T-box-tRNA | T-box-acc.11-tRNA | S3.11-Tbox-tRNA | S3.11-Tbox-acc.11-tRNA |
| --- | --- | --- | --- | --- |
| Microscope | Titan Krios | Titan Krios | Titan Krios | Titan Krios |
| Voltage (kV) | 300 | 300 | 300 | 300 |
| GIF Quantum energy filter slit width (eV) | 20 | 20 | 20 | 20 |
| Detector | Gatan K2 | Gatan K2 | Gatan K2 | Gatan K2 |
| Magnification | 130,000X | 130,000X | 130,000X | 130,000X |
| Pixel size (Å) | 1.06 | 1.06 | 1.06 | 1.06 |
| Symmetry imposed | C1 | C1 | C1 | C1 |
| Defocus range (μm) | -2.46~-3.26 | -2.80~-3.00 | -3.02~-3.68 | -2.30~-3.02 |
| Electron exposure (e-/Å2) | 49 | 49 | 49 | 49 |
| Micrographs (acquired) | 5,792 | 1,845 | 1,205 | 1,730 |
| Number of extracted particles | 3,219,341 | 632,220 | 325,698 | 623,136 |
| Number of particles after 2D classifications | 830,281 | 186,019 | 167,890 | 234,848 |
| Number of particles going to 3D refinement | 45,104 | 10,081 | 15,604 | 14,984 |
| Map resolution at 0.143 FSC criterion (Å) | 6.32 | 7.50 | 9.63 | 6.85 |
| Local resolution range (Å) | 5.81-14.04 | 7.72-12.79 | 8.41-12.43 | 8.19-16.87 |
| Sharpening B-factor (Å) | -377.6 | -311.8 | -832.8 | -289.0 |
| **Model refinement** |  |  |  |  |
| Atoms | 4,899 | 5,365 | 5,366 | 5,832 |
| Residues | 229 | 251 | 251 | 273 |
| CCmask | 0.75 | 0.75 | 0.59 | 0.80 |
| ResolutionFSC map vs. model @ 0.143 (Å) | 6.32 | 7.50 | 9.63 | 6.85 |
| **r.m.s. deviations** |  |  |  |  |
| Bond lengths (Å) | 0.004 | 0.004 | 0.004 | 0.004 |
| Bond angles (°) | 0.742 | 0.729 | 0.773 | 0.715 |
| Ramachandran favored (%) |  |  |  |  |
| Ramachandran allowed (%) |  |  |  |  |
| Ramachandran disallowed (%) |  |  |  |  |
| MolProbity score | 2.38 | 2.25 | 2.37 | 2.34 |
| Clash score | 4.61 | 3.09 | 6.00 | 4.10 |
